# The lectin Emp47/ERGIC-53 targets misfolded O-mannosylated ER proteins to ERAD

**DOI:** 10.1101/2025.11.30.691377

**Authors:** Leticia Lemus, Veit Goder

## Abstract

The yeast bifunctional O-mannosyltransferase complex Pbn1-Gpi14 generates endoplasmic reticulum (ER)-associated degradation (ERAD) signals by O-mannosylating misfolded ER proteins, but how these signals are recognized is unknown. To address this question, we identified the binding partners of Pbn1 and found that the ER lectins Emp47 (ERGIC-53 in mammals) and Emp46 were among the strongest interactors. Simultaneous deletion of Emp46 and Emp47, but not of Yos9, impaired ERAD of an O-mannosylated, N-glycan-deficient CPY* variant. ERAD was restored by plasmid-expressed Emp47, but not by a mutant lacking its carbohydrate-recognition domain (CRD). Furthermore, we show that Emp47 interacts physically with the Hrd1 complex. Together, our data support a model in which Emp47 both recognizes O-mannoses on misfolded proteins and targets them to ERAD.

## Introduction

Most proteins that enter the secretory pathway undergo N-glycosylation in the endoplasmic reticulum (ER) lumen, crucial for ER quality control (ERQC (Hammond et al., 1994; Helenius and Aebi, 2004; Caramelo et al., 2004; Molinari et al., 2004). Enzymatic trimming of N-glycans is coupled with protein folding, generating N-glycan structures that either promote (re-)binding to lectin-type chaperones for folding or, in case of terminally misfolded species, target them for delivery to ER-associated degradation (ERAD). Delivery of N-glycosylated substrates to ERAD depends on the lectin Yos9 in yeast and its homologs OS-9 and XTP3-B in mammals, which bind exposed terminal mannoses on trimmed N-glycans and, through association with ERAD machineries, provide a substrate-targeting function (Bhamidipati et al., 2005; Szathmary et al., 2005; Denic et al., 2006; Carvalho et al., 2006; Quan et al., 2008; Hosokawa et al., 2008; Christianson et al., 2008; Hosokawa et al., 2009; Clerc et al., 2009; van der Goot et al., 2018).

Using the yeast *S.cerevisiae* as model system, we have recently reported that protein O-mannosylation, a second type of glycosylation involving the attachment of single mannoses primarily to serines or threonines in the ER lumen, also generates signals for ERAD (Lemus et al., in press). This activity depends on the bifunctional O-mannosyltransferase Pbn1-Gpi14 complex (PIG-X-PIG-M in mammals) which catalyzed the O-mannosylation of various soluble and membrane-bound misfolded reporters (Lemus et al., in press). Whereas interaction and subsequent O-mannosylation of misfolded proteins by canonical protein O-mannosyltransferases (PMTs/POMTs) integrates multiple functions related to ERQC, including ER retention, prevention of protein aggregation, and release from chaperone-mediated futile folding cycles (Harty et al., 2001; Nakatsukasa et al., 2004; Goder and Melero, 2011; Xu et al., 2013), our data on Pbn1–Gpi14 suggest that protein O-mannosylation serves ultimately as a signal directing misfolded proteins for disposal via ERAD (Lemus et al., in press).

A major question concerns which cellular components recognize O-mannosylated misfolded proteins for targeting to ERAD. Using a binding assay in combination with mass spectrometry, we identified the lectins Emp46 (LMAN2, LMAN2L in mammals) and Emp47 (ERGIC-53) as prominent interactors of Pbn1. Deletion of both *EMP46* and *EMP47*, but not *YOS9*, resulted in the stabilization of a misfolded, O-mannosylated model protein lacking N-glycans. Expression of Emp47, but not of a mutant lacking the mannose-binding carbohydrate recognition domain (CRD), restored protein degradation in the double deletion background. Finally, we show that Emp47 physically interacts with Usa1, a component of the Hrd1 ERAD complex, which is involved in protein retrotranslocation.

Together, our data uncover a protein O-mannosylation–dependent ERAD pathway that relies on the ER lectin Emp47.

## Results

### Degradation of the O-mannosylated ERAD substrate ΔNg-CPY* does not require the lectin Yos9

The misfolded ER luminal protein ΔNg-CPY* is a version of the canonical ERAD substrate CPY* that lacks all its four N-glycosylation sites (Knop et al., 1996). The absence of N-glycosylation triggers O-mannosylation of ΔNg-CPY* by Pbn1-Gpi14 for subsequent Hrd1-dependent ERAD (Lemus et al., in press). To test whether the canonical ERAD lectin Yos9 was involved in the recognition and targeting of O-mannosylated ΔNg-CPY* to the Hrd1 machinery, we measured protein degradation rates in *Δyos9* deletion mutant and control cells *in vivo* after blocking protein synthesis with cycloheximide (CHX), followed by Western blot analysis. FLAG-tagged ΔNg-CPY* was stabilized only in *Δhrd1* cells but not in *Δyos9* cells (Fig. 1A-C). In control experiments, HA-tagged CPY*, was degraded faster than ΔNg-CPY*, consistent with previous reports (Lemus et al., in press), and was stabilized in both *Δhrd1 and Δyos9* cells, as expected, although to different extent (Fig. 1D-F). These data show that Yos9 is dispensable for the ERAD of the O-mannosylated substrate ΔNg-CPY* and is therefore likely not recognized by the lectin.

**Figure 1.**
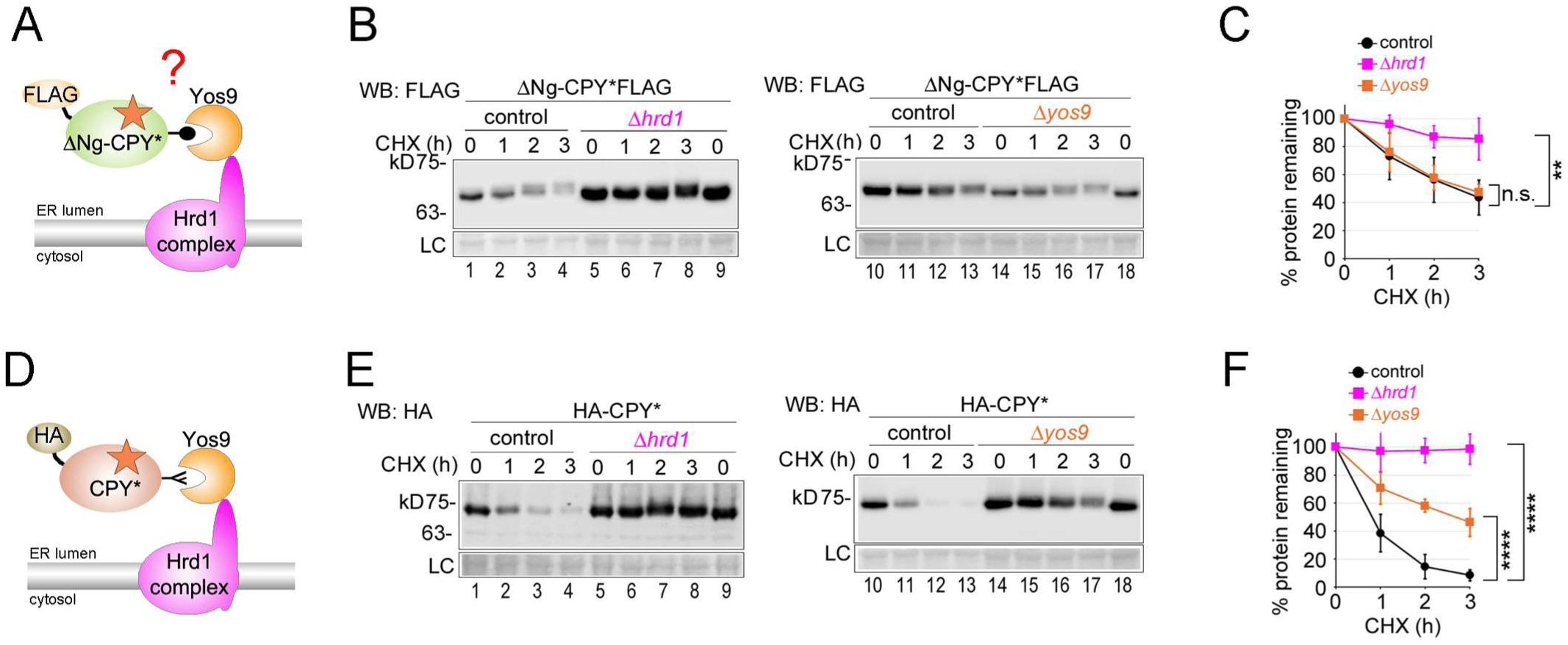
Degradation of the O-mannosylated ERAD substrate ΔNg-CPY* does not require the lectin Yos9. (**A**) Schematic depiction of the FLAG-tagged O-mannosylated ERAD substrate ΔNg-CPY*. The red star indicates misfolding. ΔNg-CPY* lacks N-glycans and the question mark indicates that it was unclear whether its mannoses (•) are recognized by the lectin Yos9, which is associated with the membrane-integrated Hrd1 complex. (**B**) Control cells and cells with the indicated deletions were used to express plasmid-borne FLAG-tagged ΔNg-CPY* for cycloheximide (CHX) shut-off experiments. Cells were lysed at indicated time points after application of CHX, and remaining ΔNg-CPY*FLAG was measured by SDS-PAGE and Western Blotting (WB) with antibodies against FLAG. Membrane staining with Ponceau served as loading control (LC). (**C**) Quantifications of results from experiments shown in (**B**). (**D**) Schematic depiction of the HA-tagged N-glycosylated ERAD substrate CPY*. The red star indicates misfolding. A critical processed N-glycan (Y) is recognized by the lectin Yos9, promoting the delivery of HA-CPY* to the Hrd1-complex for retrotranslocation into the cytosol and proteasomal degradation. (**E**) CHX shut-off experiments like in (**B**) but with cells expressing plasmid-borne HA-CPY*. (**F**) Quantifications of results from experiments shown in (**E**). Data information: Number of experiments (n) signifies biological replicates. Error bars in graphs represent standard deviation from mean. Statistical significance for all experiments involving degradation rates was calculated using two-way ANOVA (alpha = 0.05) with Šidák’s correction for multiple comparisons, obtaining the following p-values. (C): control: n=10, *Δhrd1*: n=4, *Δyos9*: n=4; p(control, *Δhrd1)*=0.0015; p(control, *Δyos9)*=0.9828; (F): control: n=10, *Δhrd1*: n=4, *Δyos9*: n=8; p(control, *Δhrd1)*<0.0001; p(control, *Δyos9)*<0.0001; n.s.=not significant; **=p<0.01; ****=p<0.0001.

### Identification of physical binding partners of the O-mannosyltransferase component Pbn1

Since O-mannosylation of ΔNg-CPY* and other substrates by the Pbn1-Gpi14 complex increased their ERAD efficiency (Lemus et al., in press), we assumed that ER-localized lectins other than Yos9 would be involved in their recognition. To find candidate lectins, we searched for interaction partners of the Pbn1-Gp14 O-mannosyltransferase complex. To this end, we N-terminally tagged Pbn1 with GFP and expressed it from its genomic locus under the control of the moderate *NOP1* promoter. Expression of GFP-Pbn1 under these conditions was fully functional (suppl. Fig. S1). Control cells or cells expressing GFP-Pbn1 were lysed and lysates were incubated with GFP-Trap for the immunoprecipitation of GFP-Pbn1, and eluted proteins were identified by mass spectrometry analysis (Fig. 2A). Among the most prominent binding partners were the lectins Emp46 and Emp47, the Emp47-associated component Ssp120, the AAA ATPase Cdc48, and the ER chaperone Kar2 (Fig. 2B; suppl. Table S1). In agreement with the mass spectrometry data indicating a physical interaction between Pbn1 and Emp47, live cell fluorescence microscopy showed that GFP-Pbn1, known to accumulate in ER puncta, partially co-localized with tdimer-tagged Emp47 (Fig. 2C and (Lemus et al., in press)).

**Figure 2.**
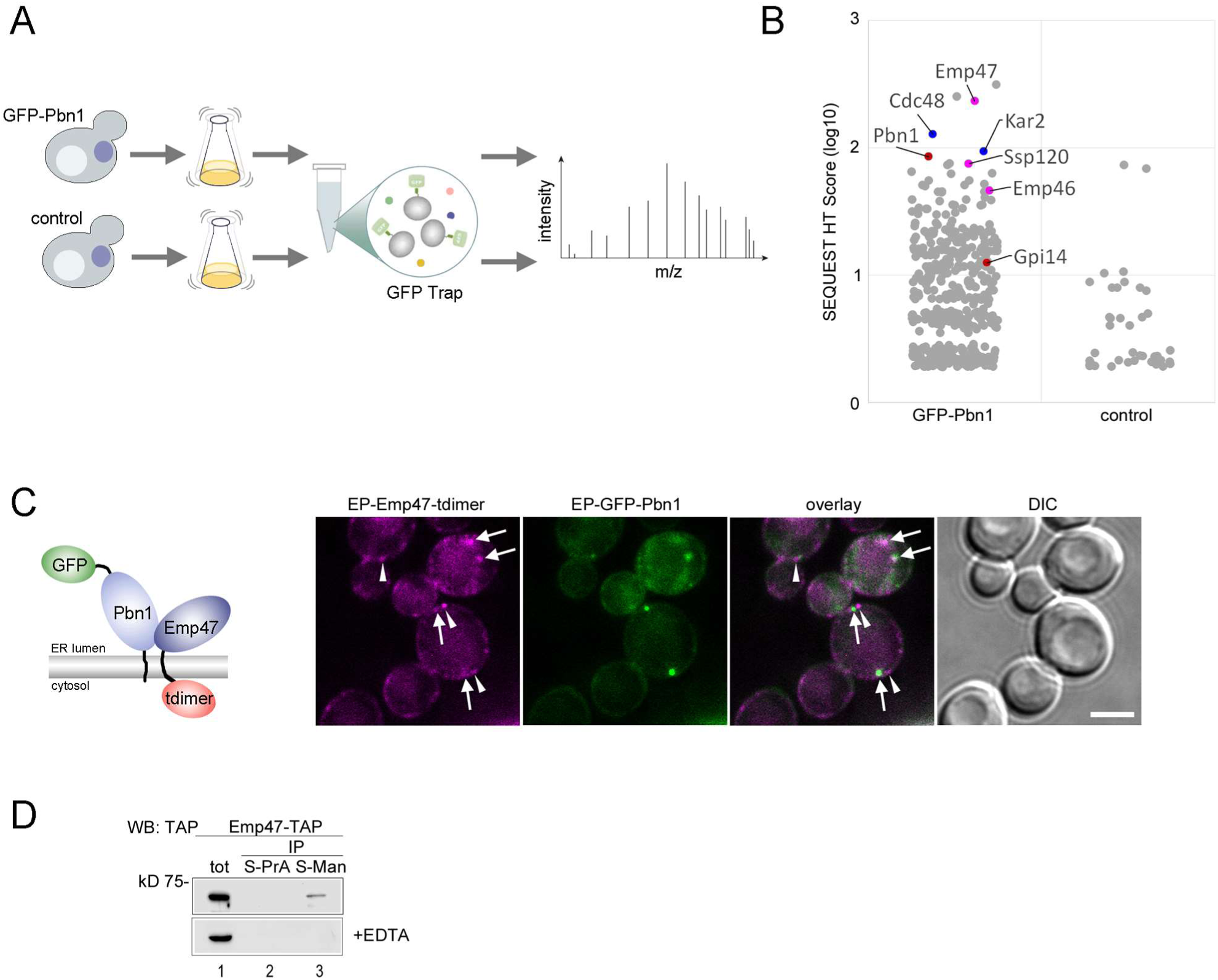
Identification of physical binding partners of the O-mannosyltransferase component Pbn1 and subsequent analyses. (**A**) Schematic illustration of the experimental approach. Cells expressing GFP-Pbn1 and control cells were grown in large cultures, followed by cell lysis and membrane solubilization in mild detergent. The cleared lysates were incubated with GFP-Trap and the eluates were analyzed by mass spectrometry. (**B**) Graphical depiction of the mass spectrometry results after following the experimental procedure described in (**A**) using cells expressing GFP-Pbn1 or control cells. Shown is a jitter plot with the SEQUEST HT score as critical parameter. Individual hits of interest are indicated. A list containing all hits is found in supplementary Table S1. (**C**) Cells co-expressing GFP-Pbn1 and Emp47-tdimer from their genomic loci and endogenous promoter (graphical depiction) were analyzed by live cell fluorescence microscopy and differential interference contrast (DIC) microscopy. Arrows indicate overlapping fluorescence signals, arrow heads indicate adjacent fluorescence signals. Scale bar: 3μm. (**D**) Mannose-binding assay. Lysates from cells expressing genomically TAP-tagged Emp47 were prepared in absence or presence of 5mM EDTA and incubated with either Sepharose-coupled Protein A (S-PrA) or Sepharose-coupled Mannose (S-Man). Eluates were used for Western Blot analysis against the TAP tag. A fraction of the lysate was loaded as input (tot).

Both, cytosolic Cdc48 and ER luminal Kar2 have known roles at distinct stages of ERAD (Jarosch et al., 2002; Carvalho et al., 2006; Denic et al., 2006; Hsu et al., 2011). The lectins Emp46 and Emp47 are paralogues and among their human homologs ERGIC-53, VIP36, VIPL, and ERGL, Emp47 and ERGIC-53 are the best studied. They are closely related and possess both a carbohydrate recognition domain (CRD) that binds to mannoses on N-glycosylated cargo inside the ER. ERGIC-53 shows Ca²⁺-dependency for this function, and in both cases, their oligomerization promotes traffic to the Golgi (Itin et al., 1996; Sato and Nakano, 2002; Satoh et al., 2006; Kamiya et al., 2008).

In support of its ability to bind mannose, we were able to isolate Emp47 from cell lysates using Mannose-Sepharose beads. To this end, we used cells expressing TAP-tagged Emp47 from the genome under the control of its endogenous promoter and performed pull-down experiments with Mannose-Sepharose beads, using Protein-A-Sepharose beads as a control (Fig. 2D). Emp47-TAP bound specifically to Mannose-Sepharose but not to Protein-A-Sepharose (Fig. 2D). In addition, we found that its binding to Mannose-Sepharose was inhibited in the presence of the chelating agent EDTA, indicating that this interaction depends on divalent cations (Fig. 2D). Given these properties, Emp46 and Emp47 emerged as prime candidates for the lectin responsible for targeting O-mannosylated ΔNg-CPY* to ERAD.

### The lectins Emp46 and Emp47 promote ERAD of O-mannosylated ΔNg-CPY*

To directly address a role for Emp46 and Emp47 in ERAD of O-mannosylated substrates, we performed CHX-chase experiments with ΔNg-CPY* in the double-deletion mutant *Δemp46Δemp47*, given the similarity between the two proteins and their likely redundant function (Fig. 3A). Strikingly, *Δemp46Δemp47* cells showed a strong delay in ΔNg-CPY* degradation (Fig. 3A, B). This contrasted with *Δyos9* cells, where ΔNg-CPY* was degraded as efficiently as in control cells (Fig. 3B and 1C). Notably, *Δemp46Δemp47* cells also reduced the degradation rate of CPY*, albeit to a lesser extent than *Δyos9* cells (Fig. 3C, lanes 1-9, D). We then generated the triple mutant *Δyos9Δemp46Δemp47* and found that it displayed an additive effect, indicating that Emp46 and Emp47 are also contributing to the degradation of N-glycosylated CPY* (Fig. 3C, lanes 10-18, D). Together, these findings indicate that Emp46 and/or Emp47 are involved in the ERAD of O-mannosylated misfolded proteins and that they may have a broader role in the ERAD of other glycoproteins.

**Figure 3.**
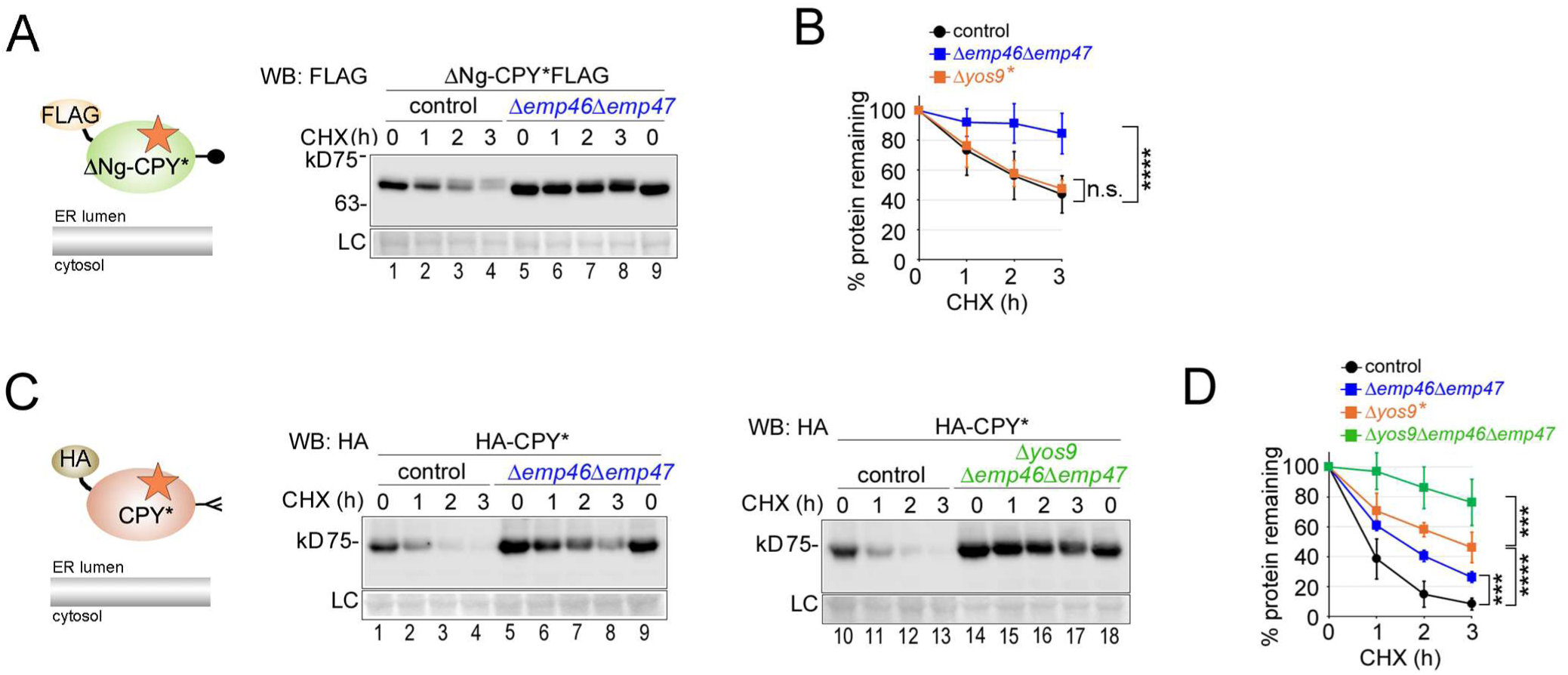
The lectins Emp46 and Emp47 promote ERAD of O-mannosylated ΔNg-CPY*. (**A**) Control cells and *Δemp46Δemp47* cells were used to express plasmid-borne FLAG-tagged ΔNg-CPY* for cycloheximide (CHX) shut-off experiments. (**B**) Quantifications of results from experiments shown in (**A**). (*) The values from experiments with *Δyos9* cells were obtained from experiments shown in Fig. 1B. (**C**) Control cells and cells with the indicated double and triple deletions were used to express plasmid-borne HA-tagged CPY* for cycloheximide (CHX) shut-off experiments. (**D**) Quantifications of results from experiments shown in (**C**). (*) The values from experiments with *Δyos9* cells were obtained from experiments shown in Fig. 1E. Data information: Number of experiments (n) signifies biological replicates. Error bars in graphs represent standard deviation from mean. Statistical significance for all experiments involving degradation rates was calculated using two-way ANOVA (alpha = 0.05) with Šidák’s correction for multiple comparisons, obtaining the following p-values. (B): control: n=10, *Δemp46Δemp47*: n=10, *Δyos9*: n=4; p(control, *Δemp46Δemp47)*<0.0001; p(control, *Δyos9)*=0.9828; (D): control: n=10, *Δemp46Δemp47*: n=4, *Δyos9*: n=8, *Δyos9Δemp46Δemp47*: n=5; p(control, *Δemp46Δemp47)=*0.0005; p(control, *Δyos9)*<0.0001; p(*Δyos9, Δyos9Δemp46Δemp47)*<0.0009; n.s.=not significant; ***=p<0.001; ****=p<0.0001.

Deletion of the gene encoding Ssp120, a known interaction partner of Emp47 involved in the ER export of glycoproteins (Margulis et al., 2016), and also identified in our mass spectrometry analysis, did not affect ERAD of either ΔNg-CPY* or CPY* (suppl. Fig. 2A-D, Fig. 2B).

### Emp47 promotes ERAD through its lectin domain and physically associates with the Hrd1-complex

Next, we tested if Emp47 function in ERAD required its CRD lectin domain. The AlphaFold structure prediction for Emp47 illustrates the boundaries and the location of its CRD. It sits N-terminally and upstream of long coiled-coil domains (stalks) which are not only involved in the formation of homo- and hetero-dimers with Emp46 but also likely provide some flexibility (Fig. 4A, and (Kamiya et al., 2008)). We generated HA-tagged plasmid-borne versions of Emp47 expressed from its endogenous promoter with and without the CRD. Both versions were expressed in comparable amounts in *Δemp46Δemp47* cells (Fig. 4B).

**Figure 4.**
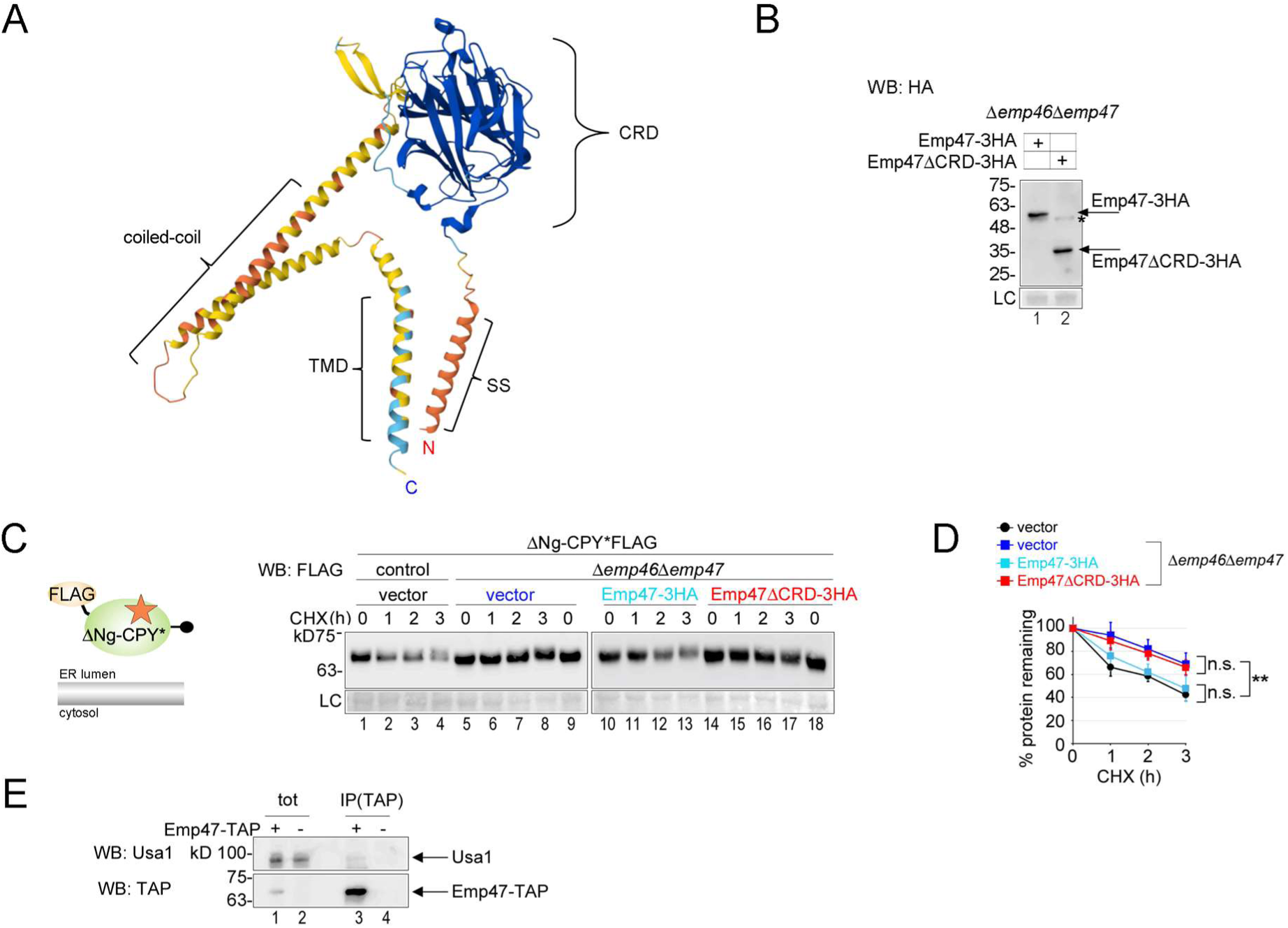
Emp47 promotes ERAD through its lectin domain and physically associates with the Hrd1-complex. (**A**) AlphaFold3 structure prediction for Emp47. The different protein domains are indicated. “SS” = cleavable signal sequence; “CRD” = carbohydrate recognition domain; “TMD” = transmembrane domain. The N- and C-terminal ends are indicated with “N” and “C”, respectively. (**B**) Plasmid-borne HA-tagged Emp47 with and without its CRD were expressed from the endogenous promoter in *Δemp46Δemp47* cells. Cell lysates were analyzed by Western Blot with antibodies against the HA tag. Membrane staining with Ponceau served as loading control (LC). The asterisk indicates an unspecific band. (**C**) Cycloheximide (CHX) shut-off experiments with control cells and *Δemp46Δemp47* cells expressing plasmid-borne FLAG-tagged ΔNg-CPY* (graphical depiction) in presence of an empty plasmid (“vector”) or plasmids expressing either Emp47-3HA or Emp47ΔCRD-3HA, as indicated. (**D**) Quantifications of results from experiments shown in (C). (**E**) Co-immunoprecipitation experiment with control cells (-) and cells expressing Emp47-TAP from the genome (+). Cells were lysed, treated with digitonin for membrane solubilization and incubated with Calmodulin-Sepharose for immunoprecipitation (IP) of Emp47-TAP. Eluates were used for Western Blot analysis with antibodies against Usa1. Emp47-TAP is recognized by the secondary antibody. Data information: Number of experiments (n) signifies biological replicates. Error bars in graphs represent standard deviation from mean. Statistical significance for all experiments involving degradation rates was calculated using two-way ANOVA (alpha = 0.05) with Šidák’s correction for multiple comparisons, obtaining the following p-values. (D): vector(control): n=3, vector: n=5, Emp47-3HA: n=6, Emp47ΔCRD-3HA: n=4; p(vector(control), vector)=0.0039; p(vector(control), Emp47-3HA)=0.6992; p(vector, Emp47ΔCRD-3HA)=0.8982; n.s.=not significant; **=p<0.01.

Next, we performed CHX-chase experiments to measure the degradation rate of ΔNg-CPY* in *Δemp46Δemp47* cells in presence of an empty plasmid or a plasmid expressing either Emp47-3HA or Emp47ΔCRD-3HA. In cells carrying an empty plasmid, *Δemp46Δemp47* mutants showed reduced ΔNg-CPY* degradation compared to control cells, as expected (Fig. 4C, lanes 1–9; D). Expression of Emp47-3HA restored ΔNg-CPY* degradation to control levels (Fig. 4C, lanes 10–13; D), whereas Emp47ΔCRD-3HA failed to do so (Fig. 4C, lanes 14–18; D). Taken together, these results show that the CRD domain Emp47 is essential for the ERAD of ΔNg-CPY*, likely for the recognition of O-mannoses on the substrate.

Finally, we asked whether we could detect a physical interaction between Emp47 and core components of the Hrd1-complex, which would be compatible with a role for Emp47 in delivering bound substrates to the ERAD machinery. To this end, we performed immunoprecipitation of Emp47-TAP and tested for binding to Usa1, a central component of the Hrd1-complex (Carvalho et al., 2006). The high molecular weight of Usa1 (90 kDa) allows for easy separation from Emp47-TAP and its potential degradation fragments, all of which are recognized by the secondary antibody but have molecular weights of 70 kDa or lower. Usa1 was visibly, albeit weakly, detectable as being co-immunoprecipitated in cells expressing Emp47-TAP, but not in control cells (Fig. 4E). These results support a model in which Emp47 engages in a transient interaction with the Hrd1 complex, thereby directly targeting O-mannosylated misfolded proteins to the ERAD machinery.

## Discussion

In this study, we identify and characterize the lectin Emp47 as a central factor in the recognition and targeting of O-mannosylated misfolded proteins for ERAD. Whereas the canonical ERAD pathway relies on N-glycan–based signals and the lectin Yos9 for substrate recognition, our data reveal a parallel, glycan-dependent degradation route in which O-linked mannoses provide the degradation signal and Emp47 acts as the corresponding reader. Our findings suggest a mechanism for how the ER interprets O-mannosylation on misfolded proteins and establish Emp47 as a functional equivalent of the N-glycan-specific lectin Yos9, but operating in an alternative, N-glycan–independent branch of ERAD.

The discovery that Emp47 recognizes O-mannosylated substrates emerged from an unbiased search for interactors of the bifunctional Pbn1–Gpi14 O-mannosyltransferase complex, which we previously showed can generate ERAD signals on both scattered and clustered serine or threonine residues of misfolded proteins (Lemus et al., in press). Among the candidates identified by mass spectrometry, Emp46 and Emp47 stood out as prominent hits, immediately suggesting a link between their capacity to bind mannoses and O-mannosylated misfolded ER proteins. The functional data presented here confirm that Emp47 is required for efficient degradation of ΔNg-CPY*, a misfolded model substrate that is O-mannosylated for ERAD. Together with the observed partial co-localization of Pbn1 and Emp47, these results suggest that O-mannosylation by Pbn1-Gpi14 and lectin-based detection by Emp47 both may occur within the same ER subdomains.

Our data further suggest that the paralogs Emp47 and Emp46 might act redundantly. In support, expression of Emp47 alone restored ΔNg-CPY* degradation to a level indistinguishable from that in control cells in a background with both lectins being deleted and with ERAD of the substrate being impaired. This rescue depended on the CRD of Emp47. The inability of Emp47ΔCRD to restore ERAD strongly supports the conclusion that the direct lectin-glycan interaction is the crucial functional step, rather than any structural or scaffolding role of the protein. This mechanism mirrors the binding of ERGIC-53 and Emp47 to high-mannose N-glycans on cargo destined for ER exit, but with a fundamentally different outcome: instead of promoting ER export of correctly folded glycoproteins, Emp47 directs misfolded O-mannosylated proteins toward degradation. How Emp47 differentiates productive cargo trafficking from misfolded-protein clearance is an intriguing open question.

One possibility is that the oligomerization states of the lectins are regulated depending on how they engage their substrates. Oligomerization of both ERGIC-53 and Emp47 has been shown to be essential for cargo export from the ER (Appenzeller et al., 1999; Sato and Nakano, 2003). Substrate binding destined for ERAD might restrict oligomerization, thereby preventing onward trafficking. Alternatively, Emp47 may simply adopt distinct oligomeric states depending on whether it functions in ERAD or in ER exit. The different types of mannose structures on N-glycans and on misfolded proteins, the latter still unexplored and potentially including variations in chain length and branching, might be important discriminating factors directing Emp47 toward ERAD or ER export. Future studies will be required to determine how Emp47 regulates these dual roles in glycoprotein handling.

A critical aspect of substrate selection for ERAD is the ability of lectins not only to bind misfolded proteins but also to physically engage the ERAD machinery. Our data show that Usa1, a core component of the Hrd1-complex, co-immunoprecipitates with Emp47, suggesting that it might directly deliver bound O-mannosylated substrates to the ERAD machinery. This is reminiscent of the interaction between Yos9 and the Hrd1-associated Hrd3 for N-glycan–dependent ERAD. Notably, previous proteomic studies have detected Emp47 associated with Yos9 and other components of the Hrd1 complex, supporting the notion that Emp47 is an integral part of a broader ERAD interaction network (Denic et al., 2006).

The additive stabilization of glycosylated CPY* observed in *Δyos9Δemp46Δemp47* cells might indicate that Emp46 and Emp47 also contribute to N-glycan-dependent ERAD. In this scenario, Yos9 possesses a high affinity for trimmed N-glycan structures, whereas Emp46/Emp47 might bind with lower affinity. Such functional redundancy or cooperativity mirrors relationships observed in mammalian systems, where ERGIC-53, VIP36 (LMAN2), and VIPL (LMAN2L) share overlapping yet distinct glycan specificities (Kamiya et al., 2008). Alternatively, apart from N-glycan processing on CPY* and ensuing recognition by Yos9, low-level O-mannosylation of the protein might also occur and could explain the reduced ERAD efficiency in cells lacking Emp47.

An intriguing observation is that Emp46 and Emp47 do not appear to be classical unfolded protein response (UPR) targets in yeast, whereas their mammalian homologs ERGIC-53 and VIP36 are transcriptionally upregulated under ER stress (Nyfeler et al., 2003). This could indicate species-specific differences for managing fluctuations in glycoprotein load. Yeast may maintain relatively constant Emp46/Emp47 levels to support both ER export and ERAD, whereas mammals may modulate lectin abundance to adapt to changing secretory or stress conditions. Whether the Emp47 function in yeast is also modulated post-translationally, through calcium binding, partner proteins such as Ssp120, or changes in subcellular distribution, remains to be studied.

Taken together, our findings define a mechanistically coherent pathway in which O-mannosylation by Pbn1-Gpi14 provides a degradation signal on misfolded proteins, Emp47 reads this signal through its mannose-specific CRD, and the Emp47–Hrd1-complex interaction ensures delivery to ERAD. This model can explain how O-mannosylation of misfolded proteins, especially those that lack N-glycans, promotes ERAD.

Future studies will be needed to define the precise structural basis of O-mannose recognition by Emp47, the regulation of its oligomeric state, and the full complement of endogenous O-mannosylated substrates that rely on Emp47 for degradation. Given the conservation of O-mannosylation and ERGIC-53-family lectins across eukaryotes, it is likely that similar mechanisms exist in mammalian cells. Identifying such pathways may have implications for diseases linked to protein misfolding, glycosylation defects, and ER storage disorders, providing new perspectives on how cells maintain proteostasis beyond the classical N-glycan–dependent ERAD paradigm.

## Abbreviations

ERQC: endoplasmic reticulum protein quality control
ERAD: endoplasmic reticulum-associated protein degradation
PMTs: protein O-mannosyltransferases
TMD: transmembrane domain
UPR: unfolded protein response

## Acknowledgements

We thank Laura Tomas Gallardo and Ana Rodriguez Hortal for invaluable help with mass spectrometry analysis. We thank Pedro Carvalho for critical reading of the manuscript and for many helpful comments. This work was supported by a grant of the Agencia Estatal de Investigación (AEI/10.13039/501100011033/PID2022-136665NB-I00) to L.L. and V.G.. L.L. was supported by the “Talento Doctor” Fellowship co-funded by the European Regional Development Fund (ERDF) and the Junta de Andalucía, Spain, and by the EMBO Scientific Exchange Grant (11267) for supporting a visiting researcher position in the Pedro Carvalho lab at the Dunn School of Pathology, University of Oxford, UK

## Author contributions

L.L. and V.G. designed and conducted experiments. evaluated data, wrote and edited the manuscript.

## Figures and legends

**Supplementary Figure S1.**
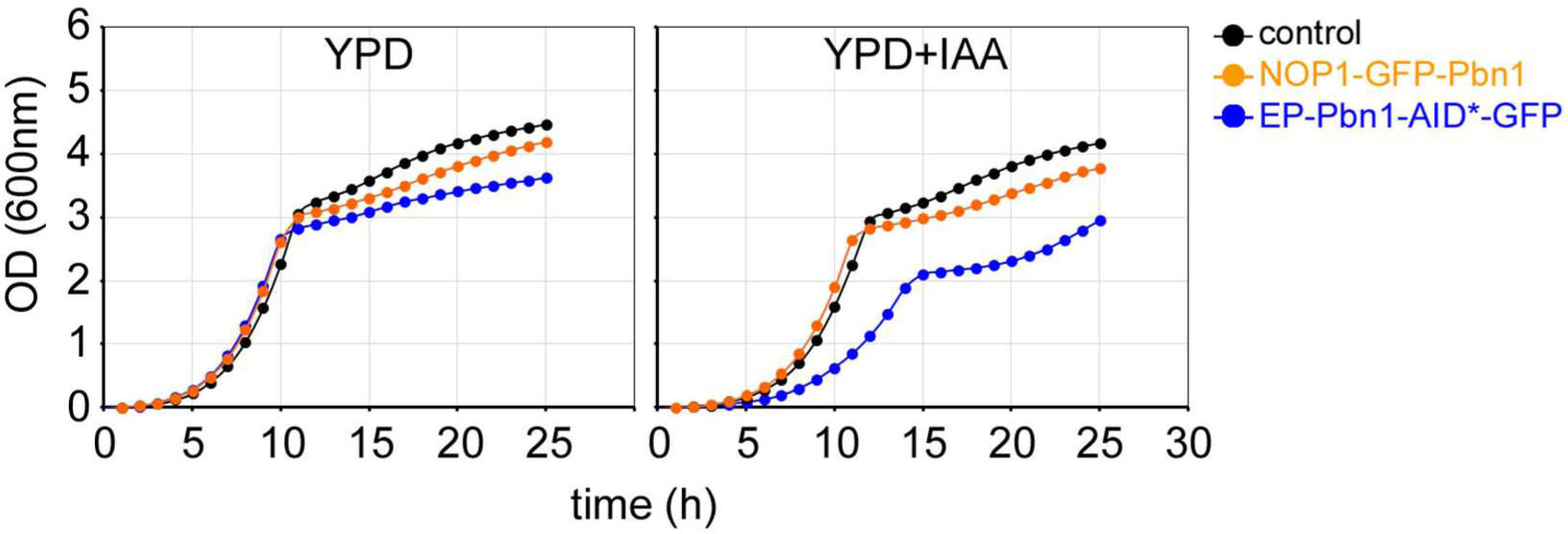
Functionality test of GFP-Pbn1 used for pulldown experiments. Wild type cells (control), cells expressing genome-integrated N-terminally GFP-tagged Pbn1 from the NOP1 promoter (NOP1P-GFP-PBN1), and cells expressing C-terminally AID*-GFP tagged Pbn1 cells expressed from the endogenous promoter (EP-PBN1-AID*-GFP), were grown in YPD medium supplemented with only solvent (YPD) or with 1mM indole-3-acetic acid in DMSO (YPD+IAA) at 30°C. Growth was determined by automated measuring of absorbance of the individual cell cultures at 600nm over a period of 25 hours. Cells expressing Pbn1 harboring the AID* degron showed a reduced growth rate in presence of IAA, due to induced degradation of essential Pbn1. Since no growth defect was seen in cells expressing NOP1P-GFP-PBN1 compared to control cells, Pbn1 is fully functional when N-terminally GFP-tagged and expressed from the NOP1 promoter.

**Supplementary Figure S2.**
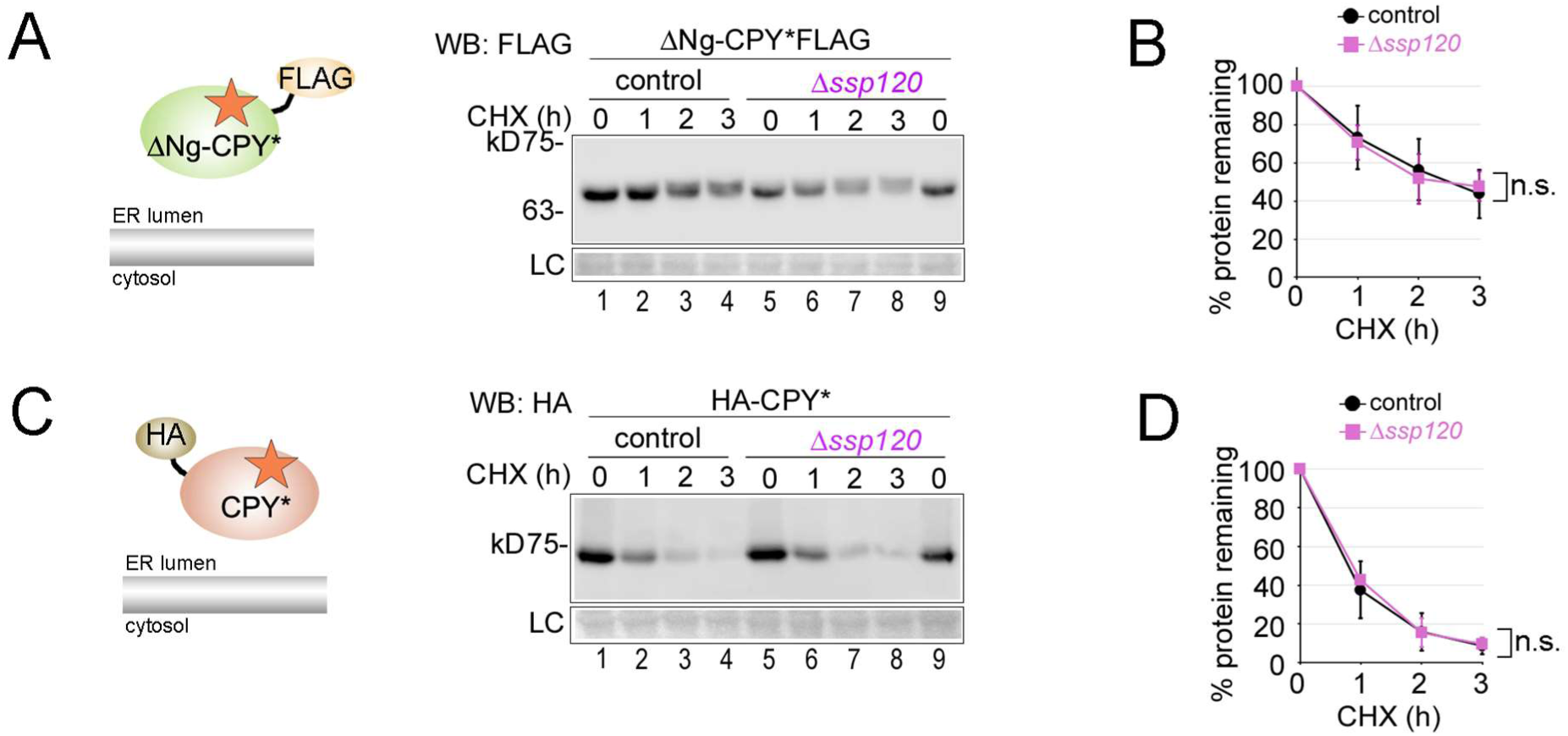
Ssp120 is not required for ERAD of either ΔNg-CPY* or CPY*. (**A**) Control cells and *Δssp120* cells were used to express plasmid-borne FLAG-tagged ΔNg-CPY* for cycloheximide (CHX) shut-off experiments. (**B**) Quantifications of results from experiments shown in (**A**). (**C**) Control cells and *Δssp120* cells were used to express plasmid-borne HA-tagged CPY* for cycloheximide (CHX) shut-off experiments. (**D**) Quantifications of results from experiments shown in (**C**). Data information: Number of experiments (n) signifies biological replicates. Error bars in graphs represent standard deviation from mean. Statistical significance for all experiments involving degradation rates was calculated using two-way ANOVA (alpha = 0.05) with Šidák’s correction for multiple comparisons, obtaining the following p-values. (B): control: n=10, *Δssp120*: n=3; p(control, *Δssp120)*=0.8905; (D): control: n=10, *Δssp120*: n=2; p(control, *Δssp120)=*0.9662; n.s.=not significant.

## Material and Methods

### Plasmids used in this study

A detailed list of plasmids used in this study can be found in supplementary Table S2. Unless stated otherwise, all constructs used in this study were expressed from centromeric plasmids. The generation of plasmids containing Emp47-3HA versions followed standard PCR-based cloning techniques using genomic DNA containing 3HA-tagged EMP47 as template. Deletion of the CRD of Emp47 was done after cloning of Emp47-3HA using the QuikChange protocol (Thermo Fisher Scientific). All relevant constructs have been verified by sequencing. All newly generated plasmids are available upon request.

### Yeast strains used in this study

All experiments were performed using common laboratory yeast strains constructed in a BY (MATa/α his3Δ1/his3Δ1 leu2Δ0/leu2Δ0 LYS2/lys2Δ0 met15Δ0/MET15 ura3Δ0/ura3Δ0) background. Genomic tagging of yeast strains was performed using standard PCR-based amplifications from suitable plasmids followed by transformation with PCR products and PCR-based verifications and/or Western Blot analysis. A detailed list of yeast strains used in this study can be found in supplementary Table S3. All newly generated yeast strains are available upon request.

### Cycloheximide (CHX) shut-off experiments

The experiments started with exponentially growing cells at 30°C in a synthetic medium with an absorbance at 600nm between 0.5 and 0.8. Translation was stopped by addition of CHX to a final concentration of 200μg/ml. Equal volume aliquots of cell culture were removed at indicated time points and moved to ice. Cells were pelleted, resuspended in 500μl cold 150mM NaOH and left on ice for five minutes, centrifuged and lysed by adding sample buffer containing 2% SDS and heating to 65°C for 10 min.

### SDS-PAGE and Western Blotting

Samples were analyzed by standard SDS-PAGE followed by Western blotting using the indicated primary antibodies, peroxidase-coupled secondary antibodies (Roche) and Enhanced Chemiluminescence (ECL, Thermo Fisher Scientific) as substrate. Images were taken with a LICOR imaging system and bands were quantified using Image Studio software (LICOR). For time course experiments, significance was calculated using two-way ANOVA with Šidák’s correction for multiple comparisons. All statistical analyses were performed in GraphPad Prism 10 software.

### Protein precipitation with Mannose-Sepharose

Exponentially growing cells were harvested by centrifugation, washed in phosphate-buffered saline (PBS, pH 7.4), and placed on ice. The cells were then resuspended in 1 mL of cold PBS containing 5 mM CaCl₂ or 5 mM EDTA and a protease inhibitor cocktail (Roche), and lysed by bead beating. The cell debris was removed by centrifugation, and the supernatant was transferred to a new tube, then supplemented with Triton X-100 (SIGMA) to a final concentration of 1% (v/v). The sample was incubated for 45 minutes with rotation at 4°C for membrane solubilization. Non-solubilized material was pelleted by centrifugation at 15,000 × g for 30 minutes, and the supernatant was carefully transferred to a new tube. An aliquot of the supernatant was removed as the input material. Mannose-Sepharose beads (Bioworld) and Protein A-Sepharose beads (Amersham) were pre-incubated with 1 mg/mL bovine serum albumin (NEB) for 30 minutes at 4°C to block nonspecific binding. Blocked beads were then added to the solubilized cell material, which had been split equally, and incubated with rotation at 4°C for 4 hours. The beads were washed four times with PBS containing 5 mM CaCl or 5mM EDTA, 1% Triton X-100, and protease inhibitors. Finally, bound material was eluted by incubating beads with sample buffer containing 2% SDS at 65°C.

### Protein co-immunoprecipitation

Exponentially growing cells were harvested by centrifugation, washed in phosphate-buffered saline (PBS, pH 7.4), and placed on ice. The cells were then resuspended in 1 mL of cold PBS containing a protease inhibitor cocktail (Roche) and lysed by bead beating. The cell debris was removed by centrifugation, and the supernatant was transferred to a new tube, then supplemented with Digitonin (SIGMA) to a final concentration of 1% (w/v). The sample was incubated for 30 minutes with rotation at 4°C for membrane solubilization. Non-solubilized material was pelleted by centrifugation at 100,000 × g for 20 minutes, and the supernatant was carefully transferred to a new tube. An aliquot of the supernatant was removed as the input material.

For the elucidation of binding partners to tandem affinity purification (TAP) tagged proteins, Calmodulin-Sepharose beads (Amersham) were used. Beads were pre-incubated with 1 mg/mL bovine serum albumin (NEB) for 30 minutes at 4°C to block nonspecific binding. Blocked beads were then added to the solubilized cell material and incubated with rotation at 4°C for 4 hours. The beads were washed four times with PBS containing, 0.2% Digitonin, and protease inhibitors. Finally, bound material, which included TAP-tagged proteins and their binding partners, was eluted by incubating beads with sample buffer containing 2% SDS at 65°C.

For the identification of binding partners to green fluorescence protein (GFP)-tagged proteins, GFP-trap (Cromotek) was used. Beads were added to the solubilized cell material and incubated with rotation at 4°C for 3 hours. The beads were washed four times with PBS containing, 0.2% Digitonin, and protease inhibitors. Finally, bound material, which included GFP-tagged proteins and their binding partners, was eluted by incubating beads with sample buffer containing 2% SDS at 65°C.

### Mass Spectrometry Analysis

The eluted fractions from the pull down were precipitated with methanol chloroform and protein pellets were resuspended in 6M urea, 50mM ammonium bicarbonate (AB). Disulfide bonds were reduced by adding DTT 10Mm (50mM, AB) to the protein solution and incubation for 60 minutes at room temperature. For carbamidometylation of cysteine –SH groups, IAA 30 mM (50mM, AB) was added to samples, followed by incubation in the dark for 30 minutes. Samples were digested overnight at 37°C using trypsin bovine (Sequencing Grade Modified Trypsin, Promega) in a ratio 1:12 enzyme-substrate. Reaction was stopped using formic acid to 0.5%. OMIX C18 tips (Agilent Technologies) were used for concentrating and desalting peptide extracts. Samples were dried, diluted in 0.1% formic acid and injected in nano-HPLC system. Protein digested samples were separated in a Thermo ScientificTM Easy nLC system using a 50cm C18 Thermo Scientific™ EASY-Spray™ column. The following solvents were employed as mobile phases: Water 0.1% Formic Acid (phase A) and Acetonitrile, 20% H2O, 0.1% Formic Acid (phase B). Separation was achieved with an acetonitrile gradient from 10% to 35% over 120 min, 35% to 100% over 1 min, and 100% B over 5 min at a flow rate of 200 nL/min. A Thermo ScientificTM Q Exactive™ Plus Orbitrap™ mass spectrometer was used for acquiring the top 10 MS/MS spectra in DDA mode. LC-MS data were analyzed using the SEQUEST® HT search engine in Thermo Scientific™ Proteome Discoverer™ 2.2 software using static carbamidomethylation (C), dynamic oxidation (M) and acetylation (C, H, K, S, T, Y) modifications. Data were searched against the Uniprot Saccharomyces cerevisiae protein database and results were filtered using a 1% protein FDR threshold. A detailed list of hits is found in supplementary Table S1.

### Growth assays

Cells were grown in the indicated liquid medium overnight at 30°C, reset to an optical density (OD600nm) of 0.05 in a volume of 500μl in a 24-well plate in duplicates and incubated for 24 hours in a BMG FLUOstar Omega multi-mode microplate reader at 700 rpm and a temperature of 30°C. ODs were recorded automatically every hour.

### Fluorescence microscopy

Exponentially growing cells were washed with PBS and immediately analyzed by fluorescence microscopy. Cells were observed with a LEICA DMi8 microscope equipped with a 100x/1.4 oil Plan-Apo immersion lens and a DIC prism and polarizer for Nomarski imaging. Images were acquired using a Hamamatsu C1 3440-20CV camera and the LASX controller software (Leica).

### Structure predictions

Protein structure prediction was performed using AlphaFold3.

